# Effects of inactivation method on SARS-CoV-2 virion protein and structure

**DOI:** 10.1101/2020.11.14.383026

**Authors:** Emma K. Loveday, Kyle S. Hain, Irina Kochetkova, Jodi F. Hedges, Amanda Robison, Deann T. Snyder, Susan K. Brumfield, Mark J. Young, Mark A. Jutila, Connie B. Chang, Matthew P. Taylor

## Abstract

The risk posed by Severe Acute Respiratory Syndrome Coronavirus −2 (SARS-CoV-2) dictates that live-virus research is conducted in a biosafety level 3 (BSL3) facility. Working with SARS-CoV-2 at lower biosafety levels can expedite research yet requires the virus to be fully inactivated. In this study, we validated and compared two protocols for inactivating SARS-CoV-2: heat treatment and ultraviolet irradiation. The two methods were optimized to render the virus completely incapable of infection while limiting destructive effects of inactivation. We observed that 15 minutes of incubation at 65°C completely inactivates high titer viral stocks. Complete inactivation was also achieved with minimal amounts of UV power (70,000 μJ/cm2), which is 100-fold less power than comparable studies. Once validated, the two methods were then compared for viral RNA quantification, virion purification, and antibody recognition. We observed that UV irradiation resulted in a 2-log reduction of detectable genomes compared to heat inactivation. Protein yield following virion enrichment was equivalent for all inactivation conditions, but the resulting viral proteins and virions were negatively impacted by inactivation method and time. We outline the strengths and weaknesses of each method so that investigators might choose the one which best meets their research goals.

## Introduction

The emergence of the novel SARS-CoV-2 resulted in tremendous social and economic distress. Factors affecting the outcome of SARS-CoV-2 infection remain unclear, meriting considerable research investment. Research with infectious SARS-CoV-2 must be performed under biosafety level 3 (BSL3) conditions. While these conditions excel at keeping researchers and the community safe, they lack the expediency required for responding to a global pandemic. Researching SARS-CoV-2 at a more accessible, lower biosafety level requires that the virus first be rendered non-infectious [1]. Several effective methods have been devised to inactivate SARS-CoV-2, but unfortunately many of these techniques destroy the structural and genetic properties required for effective viral research.

In this study, we validated a pair of techniques that can each effectively neutralize SARS-CoV-2. The two techniques—heat inactivation and ultraviolet (UV) irradiation—both inactivate SARS-CoV-2 with differing effects on important viral characteristics [2–4]. We discovered that heat inactivation is ideal for retaining genetic material while UV irradiation allows for purification of high-quality virions. We establish conditions for each method that completely inactivate viral infectivity and detail the detection of viral components that will facilitate a researcher’s particular experimental needs.

## Materials and Methods

### Virus and Cells

SARS-CoV-2 strain WA01 was obtained from BEI Resources (Manassas, VA). E6 Vero cells were obtained from ATCC (Manassas, VA) and grown in DMEM supplemented with 10% FBS, 1% pen-strep. Viral stocks were propagated and titered on E6 Vero cells in DMEM supplemented with 2% FBS and 1% pen-strep. Viral stocks were made by collecting media from infected cell cultures showing extensive cytopathic effect and centrifuged 1,000 RCF for 5 minutes to remove cellular debris. The clarified viral supernatant was then used for all subsequent inactivation studies. For determination of viral infectivity by plaque assay, E6 Vero cells were cultured then incubated with viral inoculum at limiting dilutions. Following inoculation, cells were over-layered with either 1 or 0.75% methylcellulose, DMEM supplemented with 2% FBS and 1% pen-strep and incubated for 3-4 days [1,5]. Cells were then fixed and stained with 0.5% methylene blue/70% ethanol solution. Plaques were counted and the overall titer was calculated.

### Heat Inactivation

SARS-CoV2 viral supernatants were aliquoted into 1.5 mL screw cap tubes and incubated in a 65°C water bath for defined periods of time. Triplicate samples were generated for each time point tested. The water bath was set to 65°C and validated with a thermometer. Tubes were placed into the water bath and held for 15, 20, 25, and 30 minutes. At each time interval three tubes were removed from the water bath and placed onto chilled Armor Beads to quench the inactivation. Samples were then subjected to limiting dilution and assessed by plaque assay as described above.

### UV Inactivation

To UV inactivate SARS-CoV-2 we employed a UV sterilizing oven (Fisher Cat No. 13-245-22) placed within a biosafety cabinet. This sterilizer is equipped with 5 UV bulbs with peak emission around 254 nm (UV-C irradiation). UV exposure of viral supernatants was conducted in an open top 10 cm Petri dish. Up to 15 mL of viral supernatant was placed into three separate dishes and put into the sterilizer where the lids were removed. The maximum depth of material was calculated at 1.5 mm. The dishes were irradiated for the indicated time and the power recorded. At each time point 250 uL of viral supernatant was collected from the dish. Samples were then subjected to limiting dilution as indicated and assessed by plaque assay as described above.

### RNA quantification

The sequences of qPCR amplification primers for the SARS-CoV-2 RdRp (Orf1ab) gene were: D2-8F_nCoV_RdRP forward primer 5’-GTGARATGGTCATGTGTGGCGG −3’, D2-8R_nCoV_RdRP reverse primer 5’-CARATGTTAAASACACTATTAGCATA −3’ [6]. The sequence of RdRp gene TaqMan probe was: D2-8P2_nCoV_RdRP 5’-/FAM/ CCAGGTGGWACRTCATCMGGTGATGC /BHQ1/-3’. The sequences of qPCR amplification primers for the SARS-CoV-2 E gene were: Forward Primer: D2-7F_nCoV_E 5’-ACAGGTACGTTAATAGTTAATAGCGT-3’, Rev Primer: D2-7R_nCoV_E 5’-ATATTGCAGCAGTACGCACACA-3’ [6]. The sequence of E gene TaqMan probe was: 5’-/FAM/ ACACTAGCCATCCTTACTGCGCTTCG /BHQ1/-3’. Samples were amplified using SuperScript III Platinum One-Step qRT-PCR kit (Invitrogen 11732-020) with a final reaction volume of 10 μL. Primers and probes were ordered from Eurofins Operon and were prepared as 100 μM stocks. The working stocks of the primers were 25 μM with a final reaction concentration of 800 nM. The working stock of the probe was 10 μM with a final reaction concentration of 200 nM. Each reaction mix contained 0.05 μM ROX reference dye, 0.32 U/μL SUPERase RNase Inhibitor (Invitrogen AM2694), and 1 μL of RNA. Thermocycling was performed in a real-time qPCR machine (QuantStudio 3, Applied Biosystems): 1 cycle for 30 min at 60 °C, 1 cycle for 2 min at 95 °C, and 40 cycles between 15 s at 95 °C and 1 min at 60 °C. *In vitro transcribed RNA*. Standard curves were generated using serial dilutions of *in vitro* transcribed SARS-CoV-2 RdRp (Orf1ab) and E genes. To generate the *in vitro* transcribed RNA, gBlocks were ordered from IDT with a T7 promoter, forward and reverse primer sites, and probe sequence for the RdRp (Orf1ab) gene and E gene of SARS-Cov-2 Wuhan-Hu-1 strain. The gBlocks were *in vitro* transcribed using a MEGAscript T7 RNA Synthesis Kit (Ambion, AM1333) following manufacturers instruction and purified over a GE Illustra Sephadex G-50 NICK column (Cytiva, 17085501). RNA concentration was quantified using a NanoDrop spectrophotometer to determine the copy number per μL.

### Sucrose Cushion purification

Inactivated viral supernatant was overlayed onto a two-step sucrose density cushion previously used to purify coronavirus particles [7]. The phosphate buffered sucrose at 17% and 30% were layered on the bottom of ultracentrifuge tubes. Viral supernatants were then layered on top of the sucrose and then spun at 87,000 rcf for 2 hours in either a SW28 or SW21 rotor in a Beckman ultracentrifuge. The resulting supernatant was decanted, and each pellet was resuspended in up to 300 uL of Phosphate Buffered Saline.

### Gel Electrophoresis and Western Blotting

Protein content was analyzed using SDS-PAGE and Western blot. Briefly, resuspended virion pellets were mixed with 2x loading buffer, heated to 95°C for 5 minutes then loaded onto a 10% SDS-PAGE gel. For direct detection of protein bands, gels were stained with Coomassie Brilliant Blue. For detection of specific proteins, gel separated proteins were then transferred to PVDF membranes. Membranes were then blocked with 5% dried milk in PBS-0.1%Tween followed by incubation with SARS Coronavirus NP Monoclonal Antibody (E16C) (ThermoFisher Scientific, Catalog # MA1-7403) or SARS Coronavirus Spike polyclonal Serum (BEI Resources, Manassas, VA). Primary antibody complexes were detected with goat anti-mouse (Invitrogen) or goat anti-rabbit (Santa Cruz Biotech) HRP conjugated secondary antibodies. Reactive bands were detected by ECL reagent and exposure to autoradiography film.

### Electron Microscopy

Viral samples were negatively stained by placing 5ul of resuspended virion pellets on a 300 mesh formvar coated copper grid and left for 30 seconds. Excess liquid was then wicked off and 5ul of 2% uranyl acetate was applied. This stain was also wicked off after 30 seconds. The stained grids were viewed with a LEO 912 (Zeiss) transmission electron microscope operated at 100KV accelerating voltage. Photos were taken with a 2K X 2K Proscan camera.

### Detection of viral antigens by ELISA

Suspensions of purified virions were coated onto 96-well ELISA plates at 4ug total protein per ml in PBS (50ul per well) and incubated at 4 degrees C overnight. Excess protein was decanted and plates washed five times with 0.1% Tween-20 in PBS (wash buffer). Plates were blocked with 3% nonfat milk in wash buffer for 1 hour. Mid-titer rabbit polyclonal serum obtained from BEI Resources (Manassas, VA) was 2-fold serially diluted in 1% nonfat milk in wash buffer, applied to plates, and incubated for 2 hours. Antisera was decanted and plates were washed as above, followed by incubation for 1 hour at room temperature with a 1:3000 dilution of goat-anti-rabbit secondary antibody conjugated to HRP in wash buffer with 1% nonfat milk. The extent of antibody capture was measured by colorimetric detection following treatment with TMB substrate and acid stop solution and quantified on a Molecular Devises VERSAMax microplate reader.

## Results

### Inactivation of SARS-CoV-2 by exposure to elevated temperature

The first method of inactivation we employed was the well-established procedure of incubation at high temperature [2,8]. Viral supernatants were incubated in a water bath at 65°C for specific intervals of time. The principle of this method is that excessive heat destabilizes viral proteins and assemblies, rendering them incapable of infection.

To test this inactivation method, a time course of heat exposure was conducted in screw-cap tubes containing 1.4 mL of SARS-CoV-2 viral stocks. The water bath was pre-heated for one hour with the temperature confirmed by an external thermometer. The tubes were placed into the water bath and held for 15, 20, 25, or 30 minutes. At each time point three tubes were removed from the water bath and placed onto chilled beads to reduce temperature and prevent excessive inactivation. Samples were then subjected to limiting dilution and assessed by plaque assay, as detailed in Table 1. Our calculated titer at 0 minutes was 1.04 × 10^8^ pfu/mL. Plaque assays performed on clarified viral supernatant heated at 65°C for 15, 20, 25 and 30 minutes resulted in zero countable plaques (Table 1). This data demonstrates that a complete loss of viral infectivity was observed following heat inactivation for all time points tested after T0.

**Table 1.** Heat Inactivation

|  | *Dilution*<br>*plated* | *Plaques*<br>*counted*<br>*(each*<br>*replicate)* | *Calculated titer*<br>*(pfu/mL)* |
| --- | --- | --- | --- |
| *0 min* | 10<sup>-5</sup> | 215 | 1.04 x10<sup>8</sup> |
|  | 10<sup>-6</sup> | 13 |  |
| *15 min* | undiluted | 0,0,0 | 0 |
| *20 min* | undiluted | 0,0,0 | 0 |
| *25 min* | undiluted | 0,0,0 | 0 |
| *30 min* | undiluted | 0,0,0 | 0 |

### Inactivation of SARS-CoV-2 by exposure to UV-C irradiation

To test the inactivation ability of UV-C, a time course of UV exposure was employed to determine the minimal amount of UV irradiation required to inactivate SARS-CoV-2 [4,9]. Up to 15 mL of clarified viral supernatant was placed in a 10cm Petri dish without a lid and exposed to UV-C irradiation for various amounts of time, as described in the Materials and Methods. UV-C treated virus was then subjected to limiting dilution and assessed by plaque assay. Initial testing looked at the inactivation of SARS-CoV-2 following 15s, 30s, 45s, 1 min, 2 min, 3min, and 4 min of UV-C exposure. The calculated titer of unexposed viral stocks was 4.5 × 10^7^ pfu/mL. We observed no viral plaques at all UV-C exposure times. We therefore evaluated SARS-CoV-2 inactivation following 2s, 5s, 10s, and 15s of UV-C exposure. The titer of unexposed stocks was 1.04 × 10^8^ pfu/mL. Plaques were detected following 2s and 5s exposure, correlating to 1.39 × 10^5^ pfu/mL and 10 pfu/mL, a 3- and 7-log reduction in infectivity, respectively. Exposures of 10s or greater resulted in no detectable plaques demonstrating a complete loss of virus infectivity (Table 2, Figure 1).

**Table 2.**
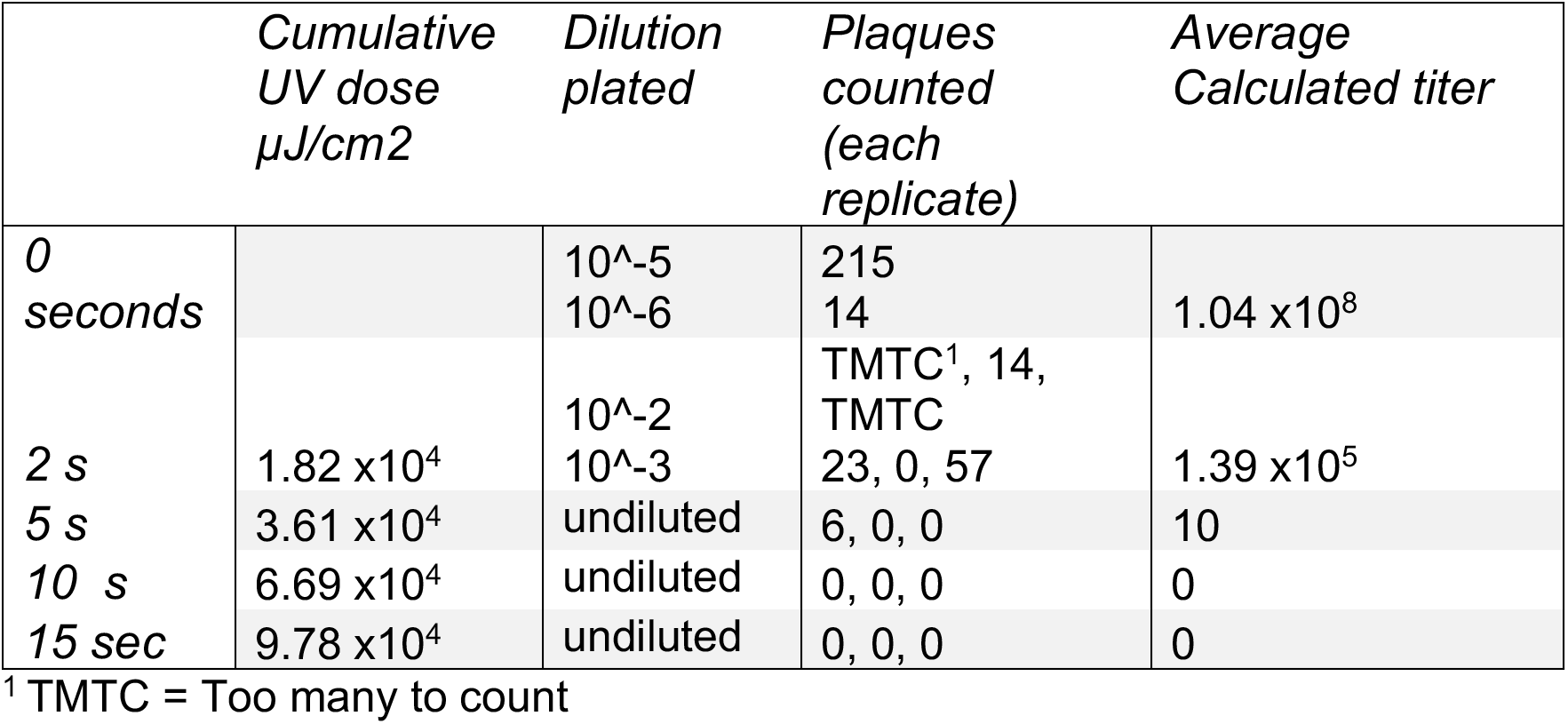
UV inactivation

**Figure 1.**
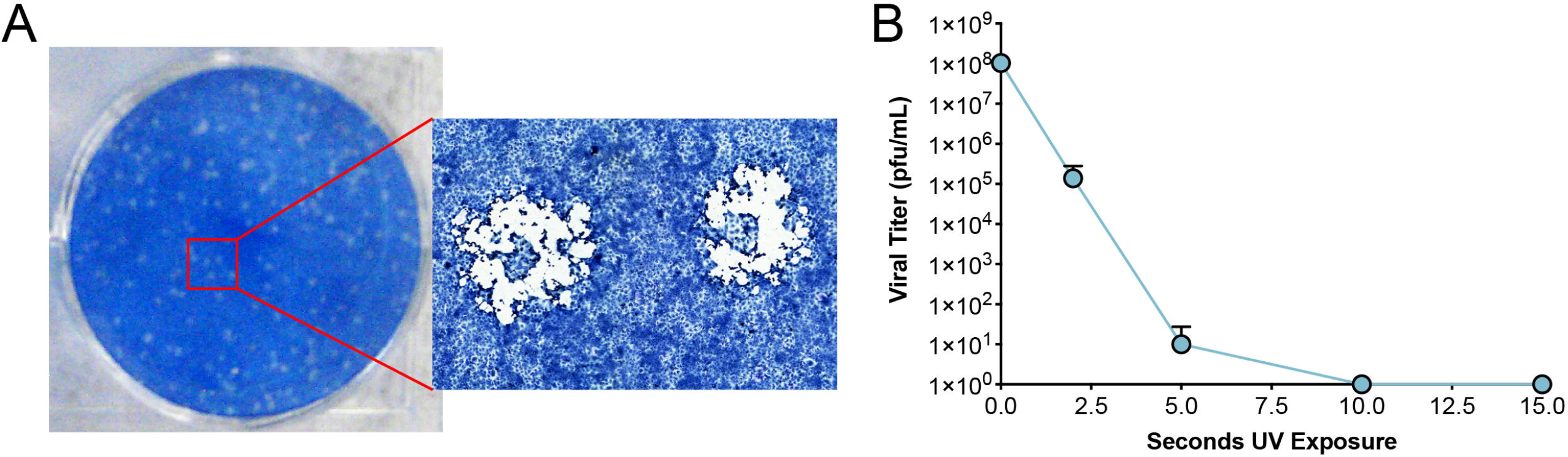
Plaque analysis and UV inactivation. A) Infectious viral particles were detected by plaque assay under methylcellulose. Depicted is a single well of a six-well plate that had been inoculated with a limiting dilution of infectious viral stock. The inset is a higher magnification image of representative SARS-CoV-2 plaques. B) A clarified solution containing infectious SARS-CoV-2 was subjected to UV exposure as detailed in Materials and Methods. Samples were taken from triplicate conditions at sequential exposures to UV-C irradiation. Each sample was then assessed for infectivity by the previously described plaque assay.

### Calculating sufficient levels of UV-C irradiation

A common method of assessing efficacy of inactivation for UV-C irradiation is to calculate a sterility assurance level [10]. The SAL is a standard used to estimate the probability of a single viable pathogen being present in a sample following inactivation. This standard is often used by manufacturers employing various irradiation-based inactivation methods to validate that products are safe. Most companies use a SAL of 10^-6^, which indicates that there is a 1 in 1,000,000 chance of a non-sterile unit surviving inactivation. For our purposes, calculating the SAL would not only help ensure inactivation of SARS-CoV-2, but also determine the minimum amount of UV-C necessary for sample inactivation. Based on intermediate inactivation values we were able to calculate the dosage of UV-C radiation that reduces infectivity of a sample by 90% or one log10 (D10 value):

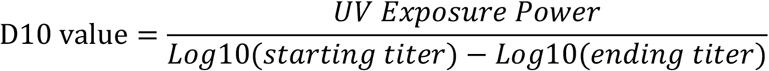

The reduction in titer following 2 and 5 seconds of UV-C exposure leads to the calculated dose needed to inactivate one log of SARS-CoV-2, which is 5,320 μJ/cm^2^ delivered by the UV sterilizer. Using this D10 value means that to reduce infectious titer of 10^8^ pfu/mL stock solution to a SAL of 10^-6^ would require an UV-C dose of 7.45 x10^4^ μJ/cm2. This is equivalent to 10.4 seconds of exposure in the UV sterilizer under the conditions we have described. It is critically important to utilize sufficient dosing of irradiation to provide a greater margin for potential error. We have chosen a minimum dose of 1.9 x10^5^ μJ/cm^2^ as sufficient to inactivate SARS-CoV-2.

### Detection of RNA from Inactivated supernatants

Various methods are currently being used to inactivate SARS-CoV-2 samples prior to diagnostic testing by quantitative Real-time PCR (qRT-PCR) or Loop Mediated Isothermal Amplification (LAMP) [11]. Given the importance of nucleic acids for these tests, we sought to determine if either of our methods would differentially impact RNA detection. We quantified the amount of RNA from both UV and heat inactivated samples and compared the number of genomes that could be detected to an untreated sample. RNA from the untreated and inactivated samples was extracted using the QiaAMP Viral RNA kit and quantified with two different primer sets targeting the RdRp (Orf1ab) or E gene of SARS-CoV-2. *In vitro* transcribed RdRp or E gene RNA was used to generate standard curves for genome copy quantification. A serial dilution of each *in vitro* transcribed RNA was performed to ensure that our different samples fell within the linear detection limit of our assay. The dilutions of each *in vitro* transcribed RNA ranged from 10^8^ to 10^3^ copies per μL and was plotted against the Ct value to determine the efficiency of each reaction. The RdRp gene primer pairs produced a reaction efficiency around 95.22% and all samples fell within the linear range of the assay (Figure 2A). For the E gene primer pairs the reaction efficiency was determined to be 98.71% and again, all samples fell within the linear range of the assay (Figure 2B).

**Figure 2.**
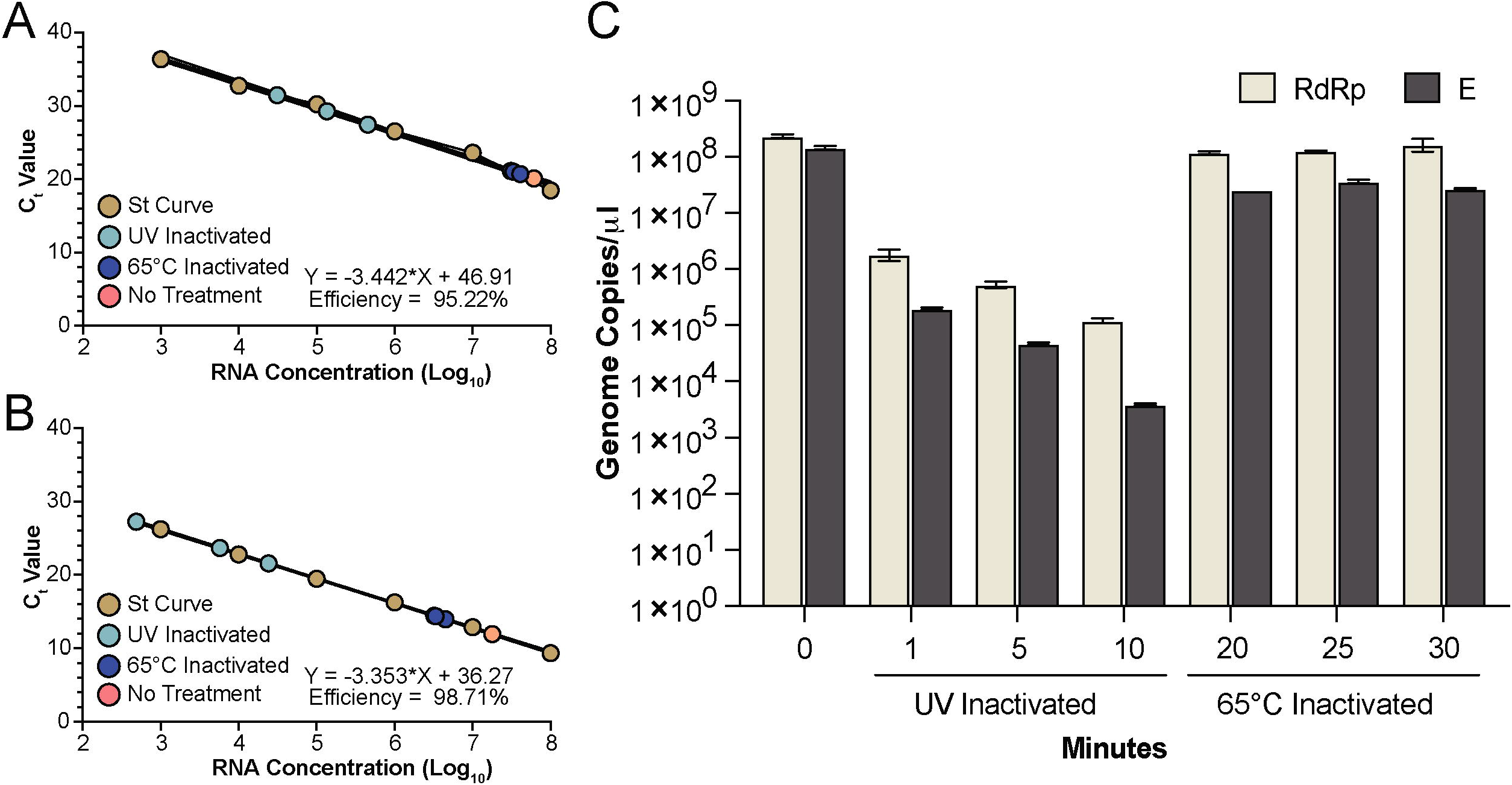
Detection of RNA genomes following inactivation. Inactivated viral supernatants were subjected to RNA extraction and detection. A and B) A standard curve was produced for the RdRp (A) and E (B) assays using serial diluted *in vitro* transcribed RNA (brown circles). RNA from UV and heat inactivated samples (light blue and dark blue, respectively) fall within the linear detection of both assays. The untreated genomic RNA is represented by the pink circle. C) Quantification of the untreated RNA, UV inactivated samples at 1, 5 and 10 minutes and heat inactivated samples at 20, 25 and 30 minutes are shown. The mean and SEM from triplicate wells for the RdRp (tan) and E (grey) assays are shown.

For the untreated sample, an average of 2.3 × 10^8^ genome copies per μl were detected by the RdRp primers and 1.4 × 10^8^ genome copies per μl were detected by the E primers. In contrast, UV inactivated samples treated for 1 minute had a 2 and 3 log reduction in the amount of RNA detected with an average of 1.8 × 10^6^ and 1.9 × 10^5^ genome copies per μl detected by the RdRp and E primers, respectively. The amount of RNA detected was further reduced as UV exposure times increased to 5 and 10 minutes. UV-C exposure for 10 minutes resulted in detection of an average of 1.2 × 10^5^ and only 3.9 × 10^3^ genome copies with the RdRp and E primers, respectively. The genomic location of the RdRp and E genes, along with the increased variability in detection via qRT-PCR, suggests that degradation of the RNA may be proceeding from the 3’ end of the genome following exposure to UV-C radiation. In contrast, heat inactivation at T20 and T25 resulted in an average of 1.2 × 10^8^ genome copies per μl and at T30 an average of 1.6 × 10^8^ genome copies per μl with the RdRp primers. The E primers detected an average of 2.6 × 10^7^, 3.6 × 10^7^, and 2.7 × 10^7^ genome copies per μl from viral stocks heat inactivated for 20, 25, and 30 minutes, respectively. Compared to UV treatment, heat inactivation did not disrupt the SARS-CoV-2 genome allowing for comparable amounts of RNA detected compared to untreated controls. Overall, the E primers resulted in slightly lower amount of RNA detected, which could be the result of degradation from the 3’ end of the genome and should be considered when designing qRT-PCR assays for use with heat inactivated samples.

### Purification of Virions from Inactivated Supernatant

To remove contaminates from inactivated virions, we employed a two-step sucrose gradient purification previously described for the porcine epidemic diarrhea virus (PEDV) [7]. Inactivated viral supernatants were subjected to high-speed centrifugation through a two-step sucrose gradient to purify and concentrate virions for downstream applications. Pelleted material was resuspended in PBS resulting in highly concentrated SARS-CoV-2 virions.

Following resuspension, we assessed the purity of viral proteins by SDS-PAGE prior to visualization by Coomassie or Western blot for viral proteins (Figure 3). Fewer Coomassie stained bands were observed from UV-C inactivated material than from heat inactivated material. This could indicate that UV-C treated virions are damaged or cross-linked together, reducing detection of SARS-CoV-2 virion proteins. More likely, the heat inactivation results in extensive protein denaturation and aggregation. These aggregated proteins may non-specifically bind to virions, or co-precipitate in our purification scheme.

**Figure 3.**
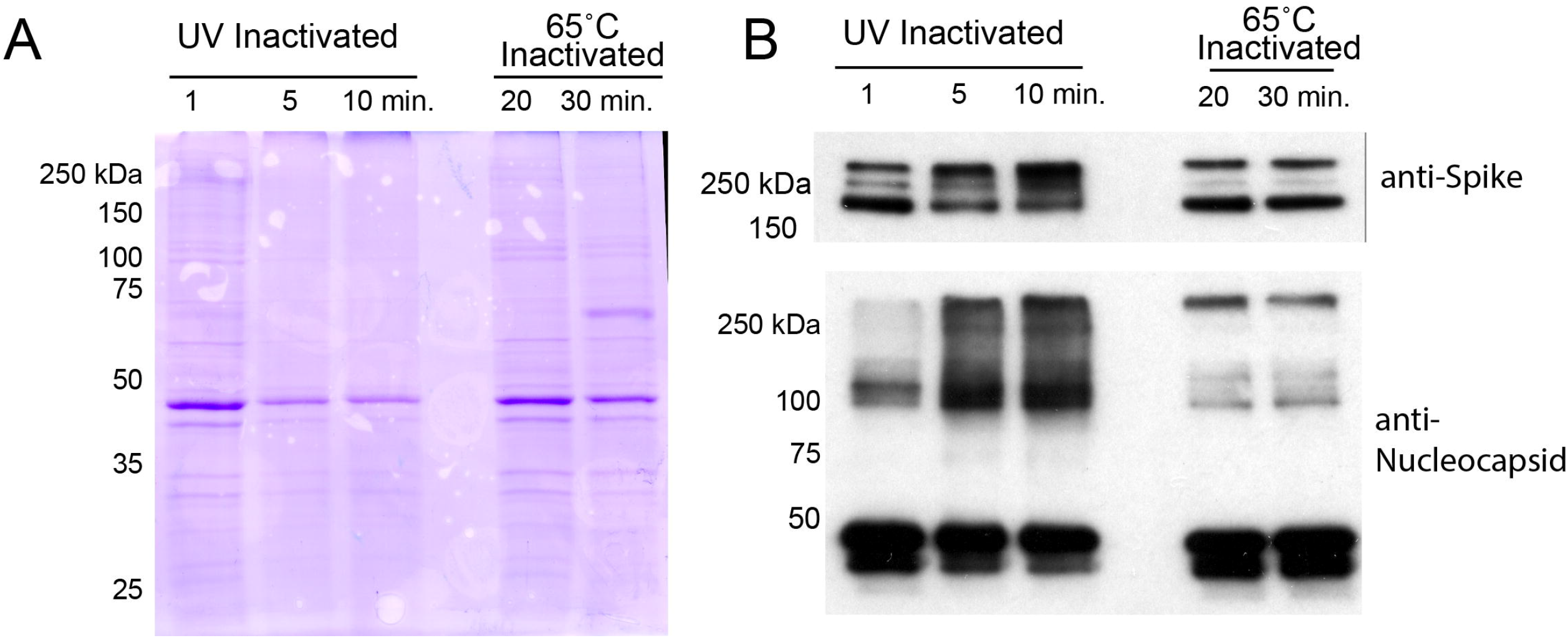
Comparison of virion protein quality following inactivation. Viral supernatants inactivated by UV or Heat exposure were loaded onto a two-step sucrose gradient and subjected to ultracentrifugation. The resulting pellets were resuspended in PBS and analyzed for protein content. A) 5 μg of resuspended pellets from the two inactivation conditions were loaded onto a 10% SDS-PAGE gel that was then subjected to Coomassie staining. B) 0.2 μg of each pellet was run on a 10% gel and transferred to PVDF membrane for Western analysis for SARS-CoV-2 Spike or Nucleocapsid. Protein extracted from SARS-CoV-2 infected cells was run as a positive control.

The differences in protein yield are also reflected in Western blot analysis of specific viral proteins. We see the most intense bands of both Spike and Nucleocapsid protein come from heat inactivated material with no obvious differences between 20 and 30 minutes of heat exposure. UV-C irradiation results in reduced quantities of both the aforementioned viral proteins. More importantly, excessive UV-C exposure at 5- and 10-minutes results in a clear increase in a slower migrating species of Spike protein. This slowly migrating species is likely the result of cross-linking between the proteolytically cleaved portions of the Spike. Additionally, we observed that greater UV exposure produced a minor, but detectable, amount of slower migrating N-protein that was not detected at 1 minute of UV exposure.

#### Electron Microscopy analysis of inactivated virions

To better understand the state of the virion following inactivation and purification, we subjected the resuspended, semi-purified virion pellets to examination under an electron microscope (Figure 4). In all conditions, spherical structures with protuberances that match descriptions of SARS-CoV-2 virions were readily detectable [12]. UV inactivated virions revealed the most intact viral particles, with the virions subjected to the lowest UV exposure appearing the most “normal”. Heat inactivated material, while mostly comparable in appearance, exhibited deformed and disrupted virion structures that increased in prevalence at longer inactivation times.

**Figure 4.**
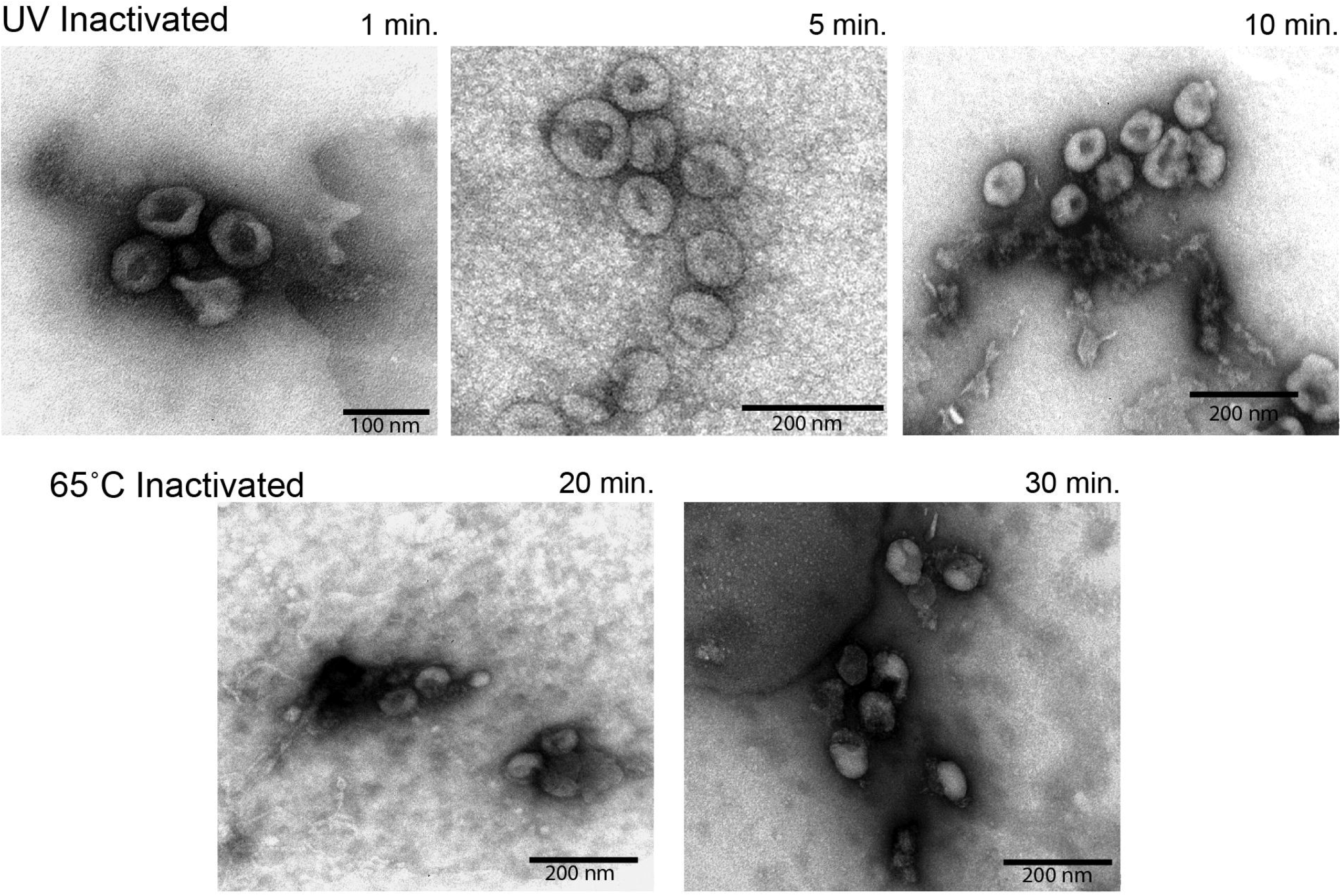
Electron microscopy analysis of virion morphology. Semi-purified virion preparations were spotted onto a grid and imaged to assess virion morphology. The top row of images was taken from UV inactivated samples. The bottom row of images was taken from heat inactivated samples. Relative size is indicated by the scale bar in the lower corner of each image.

#### Detection of inactivated and purified virions by ELISA

Retaining the antigenicity of inactivated virus is an important consideration when developing diagnostic assays or vaccines. To this end, we assessed the serological detectability of our inactivated viruses using an indirect ELISA. Wells were coated with equivalent amounts of viral protein prior to being exposed to a dilution series of SARS-CoV-2-specific poly-clonal rabbit serum. Negative control rabbit serum was used to measure the background binding capacity of the different preparations. As depicted in Figure 5, all purified virion preparations had marginal background signal from the negative control serum. We observed that detection of virion components was influenced by the type and extent of inactivation. Optimal detection was observed for samples receiving 1- or 5-minutes of UV-C irradiation, with no signal detected from the 10-minute sample. Heat inactivated material was also detected, but at a lower rate than that of UV-C inactivated samples. Together, this suggests that short duration UV exposure produces the higher quality of virions compared to heat inactivation.

**Figure 5.**
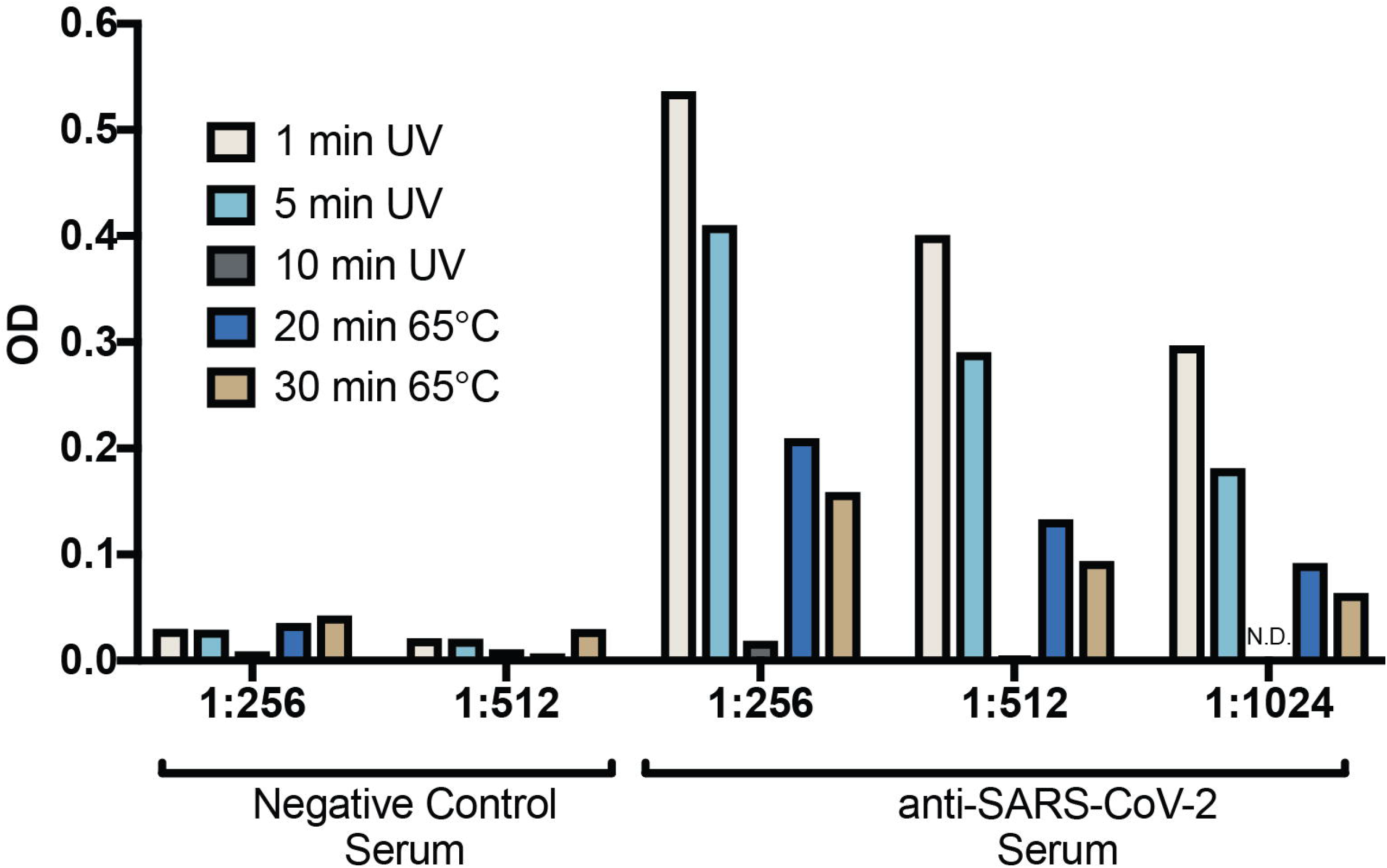
Indirect ELISA detection of virions. Semi-purified virions from both UV and Heat inactivated methods were used as coating antigen in an indirect ELISA assay. Poly-clonal serum from negative control or SARS-CoV-2 Spike immunized rabbits was used as the primary antibody with anti-rabbit-HRP and colorimetric detection was used to measure the extent of antibody capture. Plotted is the resulting O.D. measurements from a representative set of dilutions.

## Discussion

Viral inactivation is a powerful tool for mitigating research risk while expediting scientific objectives. The most useful methods of inactivation are effective and reliable without being overly destructive to virion components. The goal of this study was to validate two methods of SARS-CoV-2 inactivation—heat inactivation and UV-C irradiation—and assess their respective effects on virion components. We observed that both techniques were wholly effective at inactivating virus with minimal effects on virion morphology and antibody mediated detection.

An important initial aspect of this study was to determine the appropriate levels of viral inactivation by both methods. Insufficient inactivation puts researchers at risk for exposure, while excessive inactivation can compromise important virion components and limit research applicability. In this study, we observed that excessive UV-C irradiation reduced SARS-CoV-2 detection by ELISA, likely due to significant cross-linking of viral proteins observed by Western Blot. Many studies published on the inactivation of beta-coronaviruses use significantly higher levels of UV-C irradiation than what we report here [4,9,13]. Our methods significantly reduced UV-C exposure by optimizing experimental conditions. Working in a biosafety cabinet, we removed the plastic lids from the 10cm dishes that would have otherwise absorbed much of the incoming UV-C radiation. Because UV-C is attenuated as a function of depth, we also enhanced the surface area of exposure while limiting the fluid depth to less than 2mm, ensuring equal inactivation of the sample throughout and limiting overexposure. In a similar vein, heat inactivation was performed in small volume tubes (1.5 mL) in a water bath to ensure the even heating and inactivation of samples throughout.

It is clear that while both methods effectively inactivated SARS-CoV-2, each had unique effects on the virus that in turn affected downstream applications. Heat treatment is a common method of viral inactivation that works via the denaturation of viral proteins and disassembly of virion structures. It was interesting to see that heat inactivation, even at excessive times, left virions mostly intact, an encouraging observation for protocols that enrich virions based on the biophysical properties of intact structures. We also observed that while heat treatment eliminated infectivity, viral genomes were left largely intact. This made heat inactivation the preferred method for evaluations using genome-based assays like PCR. Unlike heat inactivation, UV-C irradiation works primarily by damaging SARS-CoV-2 RNA, preventing the transcription and replication of viral genomes. We observed this method to be especially effective at retaining virion morphology and antigenicity. Visualization of inactivated virions by electron microscopy showed that UV-C irradiated samples retained much of their native viral structure. These samples were also significantly more detectable by ELISA compared to samples that were heat inactivated. Both UV-C irradiated, and heat inactivated samples yielded near equivalent amounts of protein, though quality of viral proteins varied when assessed by Western blot. This is important to note if considering downstream applications for antigen detection or vaccine development.

The results of our study indicate that both heat inactivation and UV-C irradiation are viable methods for inactivating SARS-CoV-2 for use in BSL-2 laboratory environments. Both methods left the virion mostly intact while effects on other viral properties differed. From this study it is clear that both the extent and method of inactivation have important ramifications on SARS-CoV-2 virions that should be considered when planning experiments or downstream applications.

## Acknowledgements

The following reagents were obtained through BEI Resources, NIAID, NIH: 2019 Novel Coronavirus, strain 2019-nCoV/USA-WA1/2020, NR-52281 and Rabbit Sera Control Panels, Polyclonal Anti-SARS-CoV Spike Protein, NR-4569. Sequences for primer development were kindly provided by Dr. Dr. Jon Shultz at NIH Rocky Mountain Labs, Hamilton, MT. This work would not have been possible without the support of MSU’s Office of Research Compliance and the JRL management team of Kirk Lubick, Ryan Bartlett and Kathryn Jutila. Funding for this research was provided by the Vice President of Research and Economic Development COVID Research Fund.

